# Rigorous process for isolation of gut-derived extracellular vesicles and the effect on latent HIV

**DOI:** 10.1101/2025.01.09.632234

**Authors:** Nneoma C.J. Anyanwu, Lakmini S. Premadasa, Wasifa Naushad, Bryson C. Okeoma, Mohan Mahesh, Chioma M. Okeoma

## Abstract

**Aim:** Extracellular particles (EPs) are produced/secreted by cells from all domains of life and are present in all body fluids, brain, and gut. EPs consist of extracellular vesicles (EVs) made up of exosomes, microvesicles, and other membranous vesicles; and extracellular condensates (ECs) that are non-membranous carriers of lipid-protein-nucleic acid aggregates. The purity of EVs|ECs, which ultimately depends on the isolation method used to obtain them is critical, particularly EVs|ECs from the gastrointestinal (GI) tract that is colonized by a huge number of enteric bacteria. Therefore, identifying GI derived EVs|ECs of bacterial and host origin may serve as a window into the pathogenesis of diseases and as a potential therapeutic target.

**Methods:** Here, we describe the use of high-resolution particle purification liquid chromatography (PPLC) gradient-bead-column integrated with polyvinylpolypyrrolidone (PVPP)-mediated extraction of impurities to isolate GI-derived EPs.

**Results and Conclusion:** PVPP facilitates isolation of pure and functionally active, non-toxic EVs ColEVs from colonic contents. ColEVs are internalized by cells and they activate HIV LTR promoter. In the absence of PVPP, ColEVs have a direct reductive potential of MTT (3-(4,5-dimethylthiazol-2-yl)-2,5-diphenyltetrazolium bromide) absorbance in a cell-free system. Assessment of the origin of ColEVs reveals that they are composed of both bacteria and host particles. This protocol requires ∼12 hours (5 hours preprocessing, 7 hours isolation) to complete and should be used to purify EVs from sources contaminated with microbial agents to improve rigor. Additionally, this protocol provides a robust tool for researchers and clinicians investigating GI-derived EVs and the translational use of GI-derived EVs for diagnostic and therapeutic use.

**Highlight:**

- ColEVs but not ColECs are present in colonic content (GI tract) and can be isolated with gradient or single bead PPLC column.
- ColEVs isolated without PVPP are toxic to cells and they have a direct reductive potential of MTT. Addition of PVPP treatment in the isolation protocol results in clean and non-toxic ColEVs that transactivate the HIV LTR promoter.

## Introduction

It is now clear that EVs encompass subgroups of exosomes, microvesicles, and other membranous vesicles, with overlapping size, density, surface charge, and markers^1^. ECs, on the other hand, comprise of exomeres and supermeres that are cytosolic aggregates of lipids, proteins, and RNAs. Body fluids^2–16^, brain^17^, gut^18^ contain ECs|EVs. Cells from the three kingdoms of life––Eukarya, Bacteria, and Archaea secrete EPs, particularly EVs. Interestingly, ECs|EVs from one species may function in another species and the interspecies efficacy of ECs|EVs and their cargo has been established by our group and others^17,19^. ECs|EVs act in a paracrine fashion as regulators of health and diseases, including infectious diseases, non-communicable diseases, drug abuse, injury, reproduction, and they modulate cellular functions and/or dysfunctions^20–22^. In infectious diseases, ECs|EVs regulate viral replication and host response to infection^8,23–29^, regulate parasitic (chagas) disease^30–33^. In cancer research, EVs are explored as diagnostic biomarkers^34–38^, drug discovery, delivery, and therapeutics^39–44^. Aside from infectious diseases and cancer, EVs play key roles in the study of the pathogenesis of Asthma^45,46^, cardiovascular diseases^47^ and arthritis^48–50^, as well as in orthopedics and regenerative medicine^51–53^. The direct interaction between ECs|EVs and cells has become the basis of a short-or long-distance intercellular communication mechanism whereby ECs|EVs trigger a cellular response in host/recipient cells. Most cells respond in tissue-specific ways to ECs|EVs cargo. Thus, ECs|EVs research has become a subject of interest to scientists and clinicians across numerous disciplines from basic sciences (chemistry, biology), to applied sciences (diagnostics, pharmaceutics), and medicine. While ECs|EVs isolated from biofluids and tissues have extensively been studied, less attention has been paid to GI-derived ECs|EVs, despite the involvement and importance of the gut microbiome in health and disease, especially in the area of microbiota–gut–brain axis research. Such focus in the ECs|EVs field prompted our interest in understanding the interkingdom roles of ECs|EVs, with a focus on GI-derived ECs|EVs. However, there is a paucity of information on the effective isolation technique for GI-derived ECs|EVs.

The purity of EVs|ECs, which ultimately depends on the isolation method used to obtain them is critical, particularly EVs|ECs from the GI tract that is colonized by a huge number of enteric (pathogenic and commensal) bacteria^54^. Therefore, identifying GI derived EVs|ECs of bacterial and host origin may serve as a window into the pathogenesis of diseases and as a potential therapeutic target.

In this study, we demonstrate a rigorous protocol for isolating highly purified ECs|EVs from 1 to 2 mL colonic contents using PPLC^25,55–57^. We describe the pre-clearing of the colonic contents through flash agitation, rotation, and spinning at varying temperatures. We also describe the use of PVPP to further improve the quality of the analytes and assess the interaction of isolated analytes with host cells using functional assays.

## Materials and Methods

### Experimental design, Animal Care, Institutional/Ethical Approvals

All experiments using rhesus macaques were approved by the Tulane and LSUHSC Institutional Animal Care and Use Committee (Protocols 3574, 3581, and 3781). The Tulane National Primate Research Center (TNPRC) is an association for Assessment and Accreditation of Laboratory Animal Care International-accredited facility (AAALAC #000594). The NIH Office of Laboratory Animal Welfare assurance number for the TNPRC is A3071-01. All clinical procedures, including administration of anesthesia and analgesics, were carried out under the direction of a laboratory animal veterinarian. Animals were pre-anesthetized with ketamine hydrochloride, acepromazine, and glycopyrrolate, intubated and maintained on a mixture of isoflurane and oxygen. All possible measures were taken to minimize the discomfort of all the animals used in this study. Tulane University complies with NIH policy on animal welfare, the Animal Welfare Act, and all other applicable federal, state and local laws.

### Biospecimens

Colon content specimens used for this study were obtained from weight-matched specific-pathogen-free (free of D retrovirus, Herpes B SIV, and STLV) male Indian *Rhesus macaques* (RM, Mamu-A0*1^−^/B08^−^/B17^−^). The samples were grouped into 3: pre-infection (untreated/uninfected) and treated/infected [intravenously infected with 100 TCID_50_ dose of the CCR5 tropic SIVmac251 followed by subcutaneous administration of up to 6 MPI combination anti-retroviral therapy (ART – Dolutegravir 2.5 mg/kg, FTC or Emtricitabine 30 mg/kg, and PMPA 20 mg/kg) at 2 weeks post-SIV infection daily, and then sub-grouped into 2: one sub-group (or second group) received injections of vehicle (SIV/VEH – 1:1:18 of emulphor: alcohol: saline) two times daily and the other sub-group (or third group) received injections of Δ^9^-THC (THC/SIV) four weeks pre-SIV infection up to 6 months post-SIV infection]. Animal breeding, sample collection, and preliminary processing were carried out according to the guidelines set by the Tulane National Primate Research Center (TNPRC – AAALAC #000,594, NIH Office of Laboratory Animal Welfare assurance #A3071–01). The extracellular vesicle study was approved by the Institutional Review Board of New York Medical College (Protocol #17009). Specimens used in this study is from pre-infection (untreated/uninfected) animals.

### Cell lines

The following reagents were obtained through the NIH HIV Reagent Program has transitioned into the Biological and Emerging Infections Research Resources Program (BEI-RRP), NIAID’ s centralized research reagent repository known as BEI Resources at www.beiresources.org: TZM-GFP Human Cell Line (JC.53 Derived), HRP-20041, contributed by David G. Russell and David W. Gludish^58^; J-Lat Tat-GFP (ARP-9851) cells, contributed by Dr. Eric Verdin^59,60^. The J-Lat Tat-GFP and TZM-GFP cells were maintained in RPMI and DMEM media respectively, containing 5% EV-depleted FBS (Gibco), 100 U/ml penicillin, 100 μg/ml streptomycin, sodium pyruvate and 0.3 mg/ml L-glutamine (Invitrogen, Molecular Probes) as previously described^8^.

### General laboratory reagents

1. 1X Dulbecco’s Phosphate-buffered saline (DPBS, with calcium and magnesium, Corning, Fisher Scientific, Cat. #21-030-CM)
2. 0.1X DPBS (autoclave-sterile)
3. Polyvinylpolypyrrolidone (PVPP)
4. Sephadex beads (G-10, G-15, G-25, G-50, G-75 and G-100)
5. Sample solution (prepared as per experimental requirements)
6. #1 filter paper (Whatman, GE Healthcare, Chicago, IL)
7. 1.5% aqueous uranyl acetate (Electron Microscopy Sciences, Hatfield, PA)
8. NucBlue Live ReadyProbes Reagent (Catalog # R37606, Fisher Scientific, Waltham, MA 02451 USA)
9. DiR (1,1-dioctadecyl-3,3,3,3-tetramethylindotricarbocyanine)

### General laboratory equipment

1. Basix™ 0.2 μm Syringe Filters, Sterile (Fisher Scientific, Cat #13-100-106)
2. 15 ml Falcon tubes
3. 5 ml centrifuge tubes
4. Pipette tips 10 – 1000 µL
5. Pipettes
6. Ultrapure water from a Milli-Q IQ Water Purification System or double-distilled water.
7. Genie Vortex Mixer Model: Vortex-Genie 2
8. BioExpress Genemate Rotator W/36X1.5/2ML (with changeable paddles/rotisserie (Cat #490016-746)
9. 4°C refrigerator
10. -80°C freezer
11. Tabletop centrifuge
12. Fractionator (model)
13. Glass columns (25 ml and 100 ml, Bio-Rad)
14. Formvar and carbon-coated 400 mesh copper grids (Electron Microscopy Sciences, Hatfield, PA)
15. Glow discharge unit (e.g., PELCO easiGlow, Ted Pella, Inc., Redding, CA)
16. Parafilm (Bemis Company, Inc., Neenah, WI)
17. Transmission electron microscope (JEM-1400, JEOL, USA, Inc., Peabody, MA)
18. CCD camera (Veleta 2K x 2K, EM-SIS, Germany)
19. 24-well Costar® 24-well Clear TC-treated multiple well plate (Cat. # 3524, Corning, NY 14831 USA)
20. Greiner 96-well plate – Cat #82050-788
21. Spectrophotometer

### Development and overview of the protocol

EPs are important research, diagnostic, therapeutic, and drug delivery tools. However, despite these great attributes, challenges abound on how to isolate EPs. Currently, there are several EP isolation methods, including ultracentrifugation, density gradient, ultrafiltration, flow cytometry, immunocapture, ion exchange or size exclusion chromatography (SEC), microfluidic, asymmetric flow field flow fractionation (AF4), and precipitation^57,61^. Each of these methods has specific limitations. Some of these methods may: alter the properties of EPs; require special skills and expensive equipment; copurify all analytes with other impurities^61,62^. The hope for separating analytes, such as ECs|EVs from impurities relies in the ability to isolate and retrieve pure and functional ECs|EVs separately. This need inspired us to optimize PPLC^56,57^, which employs an innovative gradient SEC (gSEC) protocol that facilitates high-resolution separation^56^ for isolation of GI-derived EPs. Thus, in this protocol, we describe in detail the process of isolating bioactive and non-toxic ECs|EVs from colonic content of rhesus macaques. We also describe how to improve the purity of GI-derived ECs|EVs by the addition of impurity-adsorbent step using PVPP^63^. This protocol for the isolation of gut-derived EVs is modified from our published protocols where we included preprocessing sequential centrifugations and treatment with PVPP (**Figure 1A**) to the published PPLC isolation^25,26,55–57^ with fraction collection and spectral analysis with spectrophotometer (**Figure 1B**), as well as collection of different populations of analytes (ECs|EVs), aliquoting, and storage (**Figure 1C**). In previous studies, we described the use of PPLC to isolate, separate, and collect preparatory quantities of ECs and EVs from blood, semen, and tissues^25,26,55–57^, but not from intestinal contents or the gut. Below are some modifications made to PPLC, and the key steps involved in isolation of colonic ECs|EVs:

1. Soaking of Sephadex beads of various sizes (G-10, G-15, G-25, G-50, G-75 and G-100) in phosphate buffered saline at 4 °C overnight^25,55–57^.
2. Packing PPLC 1.0 × 100 cm column with the soaked Sephadex beads layered from smallest to largest^25,55–57^.
3. Pre-clearing of the colonic contents through flash agitation, rotation, spinning at varying temperatures, differential centrifugation.
4. Pretreatment of colonic contents with PVPP.
5. Loading of pre-cleared and PVPP-treated and untreated colonic contents onto PPLC column.
6. Collection of analytes with the fraction collector, spectrophotometric analysis of analyte spectral profile, and storage of analytes.
7. Downstream analysis and characterization of analytes with biochemical and cellular assays.

**Figure 1:**
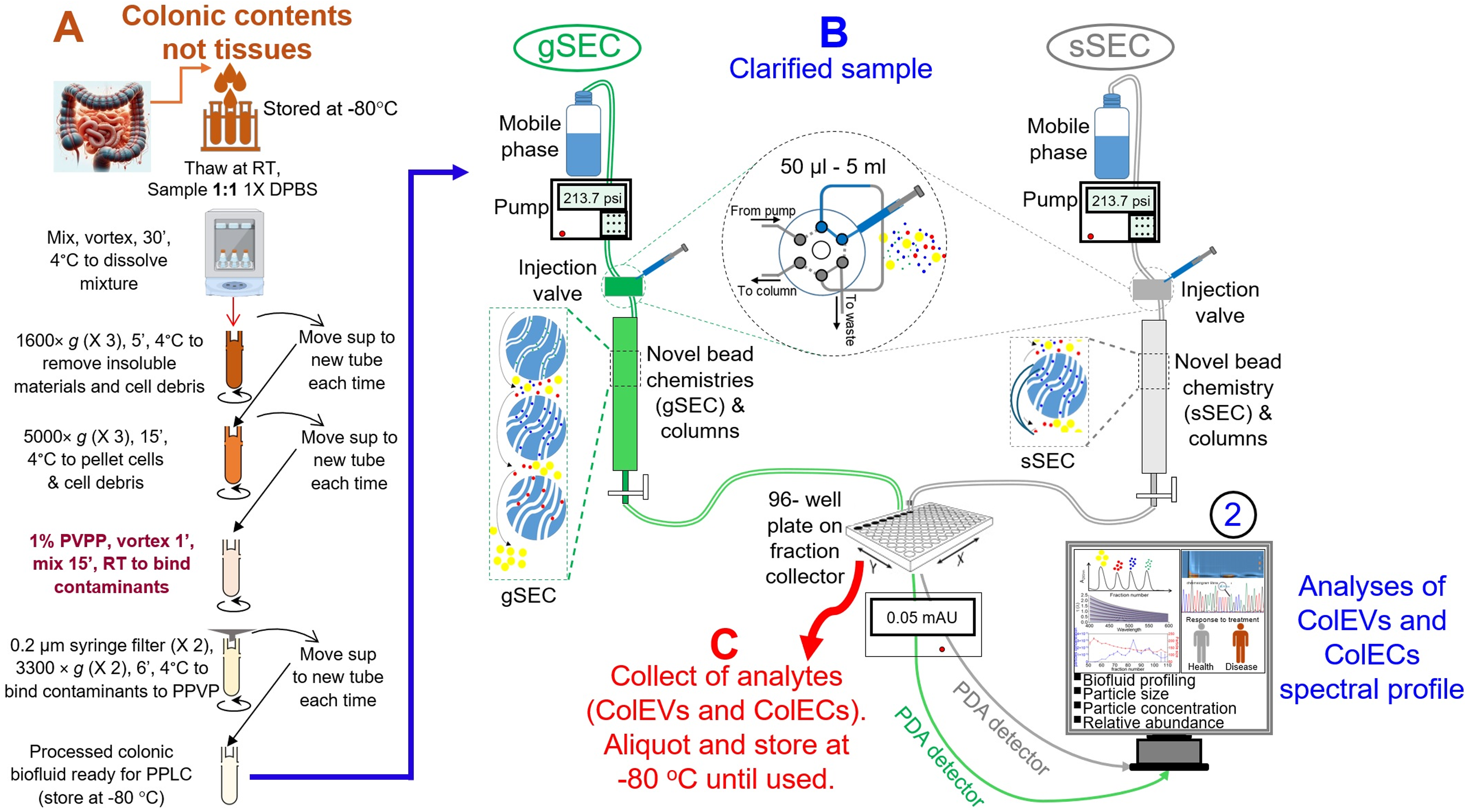
Schematic of PPLC workflow: Briefly, 300 mg tissues will be finely chopped and digested with 75 units of collagenase I at 37oC for 1 hour. The tissue digest will be clarified with minimal centrifugation steps. On-column DNAse treatment will eliminate free DNA contaminants. B) The clarified samples will be directly loaded onto gradient bead PPLC column (gSEC) or single bead PPLC column (sSEC) for ColEVs isolation. Shown in Processes gSEC, sSEC, 2 are the different aspects of PPLC all in one isolation, analysis, and retrieval steps. C) This process is independent of PPLC but represents what can be done with the analytes. This protocol is adapted from Kaddour et al., 2020 and modified for ColEVs. PVPP = Polyvinylpolypyrrolidone, gSEC = gradient bead size exclusion chromatography, sSEC = single bead size exclusion chromatography.

### Application of the protocol

ECs|EVs are gaining ground as biomarkers of health, disease, and as therapeutics tools^64–67^, for various diseases, including but not limited to, cancer, neurodegenerative disorders, and infectious diseases. The significance of ECs|EVs demands that rigorous and appropriate isolation methods, such as the use of PPLC are applied when purifying ECs|EVs from various sources, especially from complex biospecimens. The strength of this protocol is that it facilitates the use of the same starting material to separate ECs from EVs, evaluate their cargo composition, and their functions. In our prior publication on ECs|EVs, we demonstrated that ECs and EVs have distinct and overlapping cargos^25,26,56,68^. While these findings have improved our ability to isolate distinct populations of EPs, determine their composition, and biological functions. While our previous protocols have been focused of body fluid (semen, blood, urine, milk, tissue culture fluids) and tissues (brains), there is a need to develop protocol for isolation of GI-derived ECs|EVs. This present protocol provides a framework for isolation of GI-derived ECs|EVs.

The GI track is a complex environment hosting trillions of microorganisms that may exceed the number of human cells. GI-resident microorganisms secrete molecules that may affect the physiological activities of the host, including metabolism, immune response, and disease pathogenesis, such as neurological diseases^69^. Given the ability of ECs|EVs to mediate long and short distant intercellular communication, it is likely that the GI track and liver may crosstalk via ECs|EVs that delivers enteric-derived products, including toxins and GI-microbial-derived ECs|EVs to various organs, such as the liver, heart, brain to reprogram these organs. Thus, GI-derived ECs|EVs may play roles in the detection, pathogenesis, and prevention of diseases, or in the therapy of diseases.

The PPLC modified for GI-derived EPs guarantees a high yield of purified EPs that will be separated into ECs and EVs following pre-separation treatment with PVPP to remove impurities, chromatographic separation using a first-in-class gradient size exclusion column (gSEC), and collection of the different populations of EPs using a fraction collector. The collected analytes are subjected to online ultraviolet–visible (UV–Vis) monitoring of particle spectral profile. The size, yield (in concentration), and surface charge or zeta potential (ζ-potential) of the isolated ECs|EVs are assessed by nanoparticle tracking analysis (NTA), followed by transmission electron microscopy, and Western blotting analysis of EV-specific tetraspanin markers – CD9, CD63, and CD81, cellular uptake, and functional assays.

### Preprocessing of colonic contents (thawing and pre-treatment with PVPP)

All colon content samples were stored at −80 prior to processing. A starting volume of between 1 – 2 mL was thawed at room temperature in the biosafety cabinet. The following describes the sample processing or pre-isolation methods after thawing (NB: avoid freeze-thawing samples, all thawed samples should be processed before storing for isolation, if isolation is not possible on the same day).

1. On ice, collect the thawed colon contents with an RNase-/DNase-free sterile semi-microspatula (Cat #CLS3007, ThermoFisher, Pittsburgh, PA 15275, USA) and put in equal amount of 1X DPBS (1mL/gram) (Cat #MT21030CM, ThermoFisher, Pittsburgh, PA 15275, USA). For example, dissolve 1 mL of colon content in 1 mL of 1X DPBS.
2. At 4 °C, ddissolve the mixture by mixing for 30 minutes using a tube rotator (Cat #9778990, VWR, Wayne, PA 19087, USA) while vortexing (Cat #NC9864336, ThermoFisher, Pittsburgh, PA 15275, USA) intermittently on a high speed for 30 seconds every 10 minutes.
3. Remove insoluble materials and cell debris from the colon content – DPBS mix by centrifuging the mixture three (3) times at 1600 ×g at 4 °C for 5 minutes. After each centrifugation (Cat #05-400-61, ThermoFisher, Pittsburgh, PA 15275, USA), carefully collect and place supernatants into new tubes before centrifuging again.
4. Repeat centrifugation three (3) times at 5,000 × g at 4 °C for 15 minutes. Place supernatants into new tubes each time before centrifuging again.
5. Place supernatants in fresh sterile 15 mL Falcon tube and add 0.1 to 1% polyvinylpolypyrrolidone (PVPP). The PVPP binds to compounds that may interfere with the EVs isolation and improves the quality of yield.
6. Vortex for 1 minute and rotate on the tube rotator for 15 minutes at room temperature.
7. Filter into a new 15 mL Falcon tube with 0.2 µm syringe filter (Cat #13-100-106, ThermoFisher, Pittsburgh, PA 15275, USA), and centrifuge at 3300 x g for 6 minutes at 4 °C. This is to remove leftover pelleted PPVP-bound contaminants.
8. Repeat step 7. (NB: Keep samples on ice in-between steps except when room temperature is required).

Resulting colonic biofluid is ready for use for the isolation of EVs by PPLC. Samples can be stored at −80 if isolation will not be carried out the same day.

### Packing and equilibration of PPLC column for isolation

1. Make 0.1X DPBS by 1 mL of 1X DPBS in every 10 mL of double-distilled water (ddH_2_O). The 0.1X DPBS are used to soak the beads with which the PPLC columns are packed, to equilibrate the column, and to rinse the column after isolation.
2. The 100 cm gradient PPLC column was packed with multi-sized beads as previously described^56^. An empty glass column of 100 cm length, 1 cm inner diameter, and 79 ml volume (Bio-rad cat #7371091) was packed in-house with a gradient of epichlorohydrin cross-linked dextran beads of various exclusion limits controlled by different degrees of cross-linking. The beads are commercially available from Cytiva and sold under the trade name Sephadex (previously branded for GE Healthcare). The characteristics of the beads are described in the table below. The beads were slowly packed from bottom to top after overnight swelling in 0.1X DPBS, starting with G-10 and ending with G-100 (**Table 1**).
3. The column is connected to a low-pressure, drop-based small-volume fraction collector (SKU: 171041, Gilson Incorporated, Middleton, WI 53562-0027) which collects the isolated EVs in microplates for UV-Vis spectrometry.
4. With the prepared 0.1X DPBS, equilibrate the gradient PPLC column with 5 ml of PBS prior to usage. To confirm that the column is ready-to-use, run the 0.1X DPBS collected in the microplate on Biotek Synergy H1 spectrophotometer/microplate reader (Cat #11120531, ThermoFisher, Pittsburgh, PA 15275, USA). A reading less than 1 nm at 230 nm absorbance indicates that the column is well-rinsed and ready-to-use. (Different brands of microplate may give background absorbance. The recommended microplate for use is Greiner – Cat #82050-788, VWR International, Wayne, PA 19087, USA).

### Isolation of colon content fluid via PPLC

1. Place new plates on the stage of the fraction collector
2. Pipette the colon content fluid onto the gradient column and collect fractions in 96-well with fractionator.
3. Check the absorbance spectra at ranges 280 – 650 nm on spectrophotometer
4. Plot a graph of the absorbance peaks (see Fig. 2) to determine the EV-rich fractions
5. Pool fractions that are rich in EVs (EV-rich fractions form the first peak on the UV-Vis spectre).
6. Store at −80 for further downstream application.

### Notes

- Columns can be reused for as long as the flow rate is optimal. This is usually between 4^th^ – 7^th^ sample isolation.
- Ensure that microplates are washed by soaking for at least 1 hour with 5% bleach, rinsing with absolute ethanol, and finally with double distilled water before reuse.

### Nano Tracking Analysis (NTA)

Samples were diluted to the appropriate concentration in filtered 0.1 X PBS for NTA. Size distribution, particle concentration and Zeta potential (ζ potential) of the purified ColEVs were determined using ZetaView PMX110 (Particle Metrix, Mebane, NC, USA). First, the ZetaView machine and software were initialized, followed by a CellCheck to ensure the measurement cell. The cell assembly was flushed with deionized water to remove any contaminants. An alignment suspension (dilution 1:250.000) was then used to calibrate the instrument, which involved injecting the suspension, avoiding air bubbles, and allowing the instrument to perform auto-alignment and focus optimization. The system was calibrated using 100 nm Nanosphere™ size standards (3100A, Thermo Fisher Scientific) before acquisition of measurements. The instrument parameters were set up, including temperature (25 °C), shutter speed (70), camera sensitivity (92), and frame rate (30 fps). During auto-alignment, the instrument finds particles, focuses on the particles first in large steps followed by small steps and finally performs a zeta potential measurement for profile symmetrization. A symmetric and smooth profile indicated that the instrument was ready while a curve that was inverted, jagged or flat indicated air bubbles inside the cell or a cue to clean the cell. Samples were then introduced into the chamber (in sample 1:1000 0.1X DPBS dilutions) and measurements were taken in triplicates at multiple positions to ensure accuracy. The ZetaView tracked the Brownian motion of particles to determine their size and concentration, utilizing sliding optics to move between focal positions and sample a large volume of the solution. For size, automated analysis of the 11 positions was done, any outlier position was removed, and the median number (X50) was used to report the particle size. The measured concentration was normalized to the original volume of ColEVs before dilution and reported in particles/mL. Additionally, the instrument measured the zeta potential of particles by generating particle drift in an electrical field.

**Figure 2:**
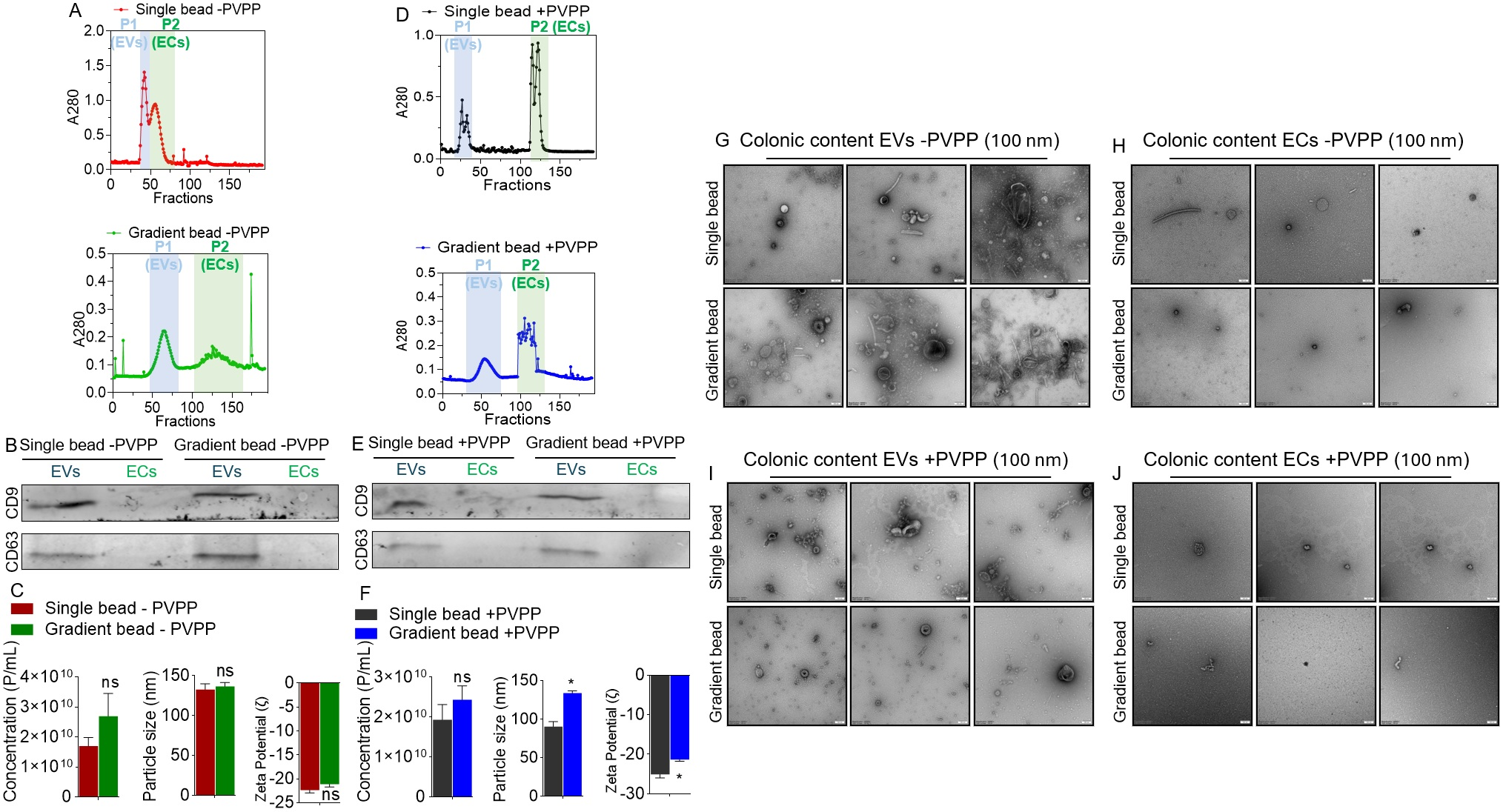
Integration of PVPP and PPLC bead-column chemistry facilitates isolation of clean ColEVs: **A**) Spectra of the analytes isolated with single or gradient beads in the absence of PVPP (-PVPP). **B**) Western blot of known EV markers present in ColEVs isolated with single or gradient beads in the absence of PVPP. **C**) Physical properties of ColEVs isolated with single or gradient beads in the absence of PVPP. D) Spectra of the analytes isolated with single or gradient beads in the presence of PVPP (+PVPP). **E**) Western blot of known EV markers present in ColEVs isolated with single or gradient beads in the presence of PVPP. **F**) Physical properties of ColEVs isolated with single or gradient beads in the presence of PVPP. **G-J**) TEM images of ColEVs isolated with single or gradient beads in the absence of PVPP (**G-H**) and in the presence of PVPP (**I-J**). P1 = Peak 1, P2 = Peak 2.

**Table 1.**
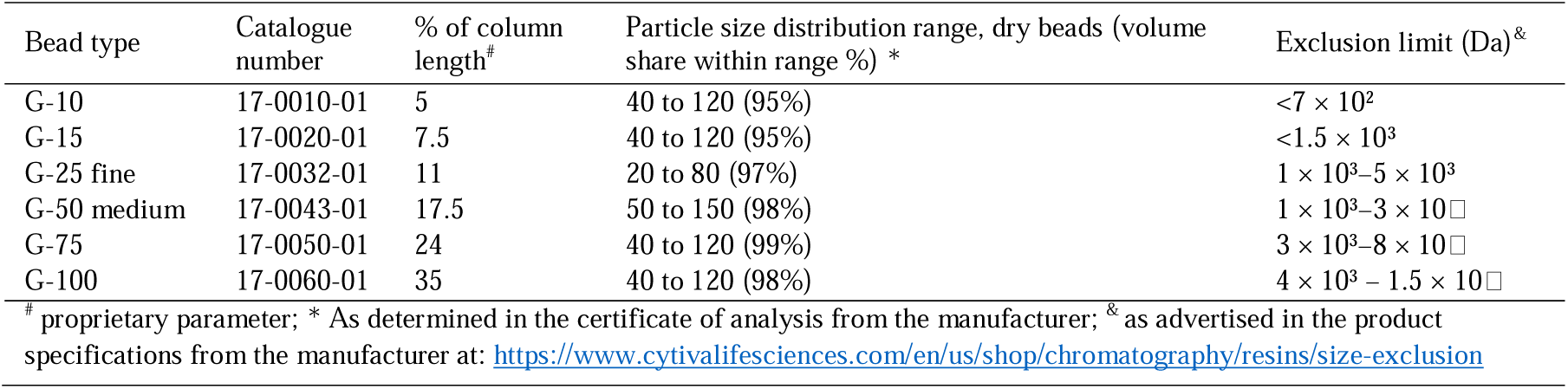
Beads characteristics.

### Labelling of EVs and uptake of EVs by TZM-GFP and JLAT-TAT-GFPcells

We determined EV uptake by TZM-GFP and JLAT-TAT-GFPcells at various incubation periods. 1ul DiR (5uM DiR-1,1-dioctadecyl-3,3,3,3-tetramethy-lindotricarbocyanine iodide - in 0.1 X PBS) was added to 100ug of EV and incubated at 37°C for 30 minutes in the dark (inverted every 5 minutes). The mix was purified in EV purification column by first, washing column with 180ul of 0.1 X PBS and running 100uL labeled EV through the column. 180ul 0.1 X PBS was then run through column, and the purified EV was eluted with 50ul 0.1 X PBS. Concentrations of labelled EV used for uptake were 0, 50, 100, 150 and 200ug each in 10,000 TZM-GFP or JLAT-TAT-GFPcells. For the JLAT-TAT-GFPcells, at 24 hours incubation, 1 drop per 5 mL of Nucblue working solution was added, 100 uL per well, and incubated at room temperature for 5 minutes before imaging. Uptake was captured and analysed at Brightfield, DAPI, GFP and Texas Red channels.

### Western blot

We also determined the presence of EV-specific tetraspanin proteins – CD63 and CD9 – by Western blotting analysis. Protein concentration of the EVs were normalized via Bradford assay using Quick Start™ Bradford BSA standards (cat #5000201, Bio-Rad Laboratories, Inc, Hercules, California 94547). 20 µg analytes representing ColEVs or ColECs were boiled at 95 °C in the presence of 4X Laemmli buffer (cat #1610747, Bio-Rad Laboratories, Inc, Hercules, California 94547) for 5 minutes before loading onto a 4–15% Tris-glycine gradient gel. The proteins were transferred onto a PVDF membrane overnight at 35V constant volt in a cold room. The membrane was then blocked with 5% BSA for 1 hour before incubating overnight (at 4°C) with primary antibodies (1:1000) against CD63, CD9 (mouse, Developmental Studies Hybridoma Bank, DSHB, Iowa City, IA, USA). After 3 washes, the blots were incubated for 2 hours at room temperature with fluorescent-labeled secondary IRDye 800CW donkey anti-mouse IgG antibodies (LI-COR, Lincoln, NE, USA), the membranes were imaged with the LI-COR Odyssey Infrared Imaging System.

### Transmission Electron Microscopy (TEM)

Formvar and carbon-coated 400 mesh copper grids (Electron Microscopy Sciences, Hatfield, PA) were glow-discharged for 30 seconds using a PELCO easiGlow™ unit (Ted Pella, Inc., Redding, CA) to render the surface hydrophilic. The sample solution was prepared by dispersing the specimen in an appropriate buffer to achieve a final concentration suitable for transmission electron microscopy (TEM) analysis. A 3-5 µL aliquot of the sample solution was applied to the glow-discharged grid and allowed to settle for 1 minute at room temperature. Excess sample was removed by gently touching the edge of the grid with #1 filter paper (Whatman, GE Healthcare, Chicago, IL) to wick away the solution by capillary action. The grid, with the sample side facing down, was immediately placed onto a 50 µL drop of 1.5% aqueous uranyl acetate (Electron Microscopy Sciences, Hatfield, PA) on a sheet of parafilm (Bemis Company, Inc., Neenah, WI). After 1 minute of staining, the uranyl acetate was wicked off using filter paper, and the grid was allowed to air dry for 5 minutes at room temperature. Samples were then viewed using a JEM-1400 transmission electron microscope (JEOL, USA, Inc., Peabody, MA) operated at 100 kV. Images were captured with a Veleta 2K x 2K CCD camera (EM-SIS, Germany).

### Activation of HIV LTR promoter

We assessed the activation of HIV LTR promoter by running a 24-hour kinetics on Lionheart FX Automated Microscope (SN: 18022029, Agilent Technologies, Santa Clara, CA 95051 USA). Ten thousand (10,000) TZM-GFP cells each were seeded into four wells of a 24-well Costar® 24-well Clear TC-treated multiple well plate (Cat. # 3524, Corning, NY 14831 USA) and incubated for 3 hours at 37 °C, 5% CO to allow for cell attachment. DiR-labeled EVs were prepared by adding 50 µg (per treatment/sample) of EVs to DiR in 0.1X PBS (1,1-dioctadecyl-3,3,3,3-tetramethylindotricarbocyanine) to reach a 5 µM final concentration. The labeled EVs were then passed through Cellgs EX03-8, Exo-spin columns mini (Cat. # EX03-8, CellGS LLC, St. Louis, MO, 63132, USA) for purification, and eluted with 0.1X PBS. Each well containing TZM-GFP cells was uniformly treated with the purified EVs (50 µg). 10 µL of NucBlue Live ReadyProbes Reagent (Catalog # R37606, Fisher Scientific, Waltham, MA 02451 USA) was also added to each well just before placing in the Lionheart FX Automated Microscope (SN: 18022029, Agilent Technologies, Santa Clara, CA 95051 USA).

Imaging was performed every hour in two batches, first for the first 13 hours, then for the last 5 hours (20^th^ to 24^th^ hour) using Gen5 3.15 software, configured for 10X magnification. The instrument temperature was set to 37 and the CO_2_ pump was turned on for consistent 5% CO_2_ inflow. Kinetics option was selected on the Gen5 software, the imaging channels were set up for GFP (Excitation 488 nm, Emission 509 nm), DiR (Texas Red, Excitation 748 nm, Emission 780 nm), and NucBlue (DAPI, Excitation 360 nm, Emission 460 nm).

Images were acquired at multiple time points to monitor the kinetics of uptake of ColEVs. For each time point, images were captured in all four channels (GFP, Texas Red, DAPI, and Brightfield) for each well. The acquired images were processed using Gen5 3.15 software, with background subtraction using the image processing option.

The processed images were analyzed to quantify uptake of ColEVs by measuring the fluorescence intensity of DiR-labeled EVs within GFP-positive cells. NucBlue staining was used to identify and analyze the nuclei, ensuring accurate cell counting. Using GraphPad Prism, the normalized fluorescence intensity of the cells for each channel was plotted against time to generate a kinetic uptake curve. Data were normalized to the number of cells to account for variations in cell number across wells.

### Normalization approach for cell-based assays

We divided uptake or LTR values with NucBlue values for each image. Then, multiply by initial cell number (10,000) and add a pseudo-count (1).

## Results

### EVs but not ECs are present in colonic contents of rhesus macaques

Clarified supernatants from colonic contents obtained from RMs before infection and/or treatment (pretreatment), treated with or without PVPP were loaded a top 1) single-bead or 2) gradient-bead PPLC beds^56^ to i) gain insight into the spectra of colonic contents obtained through the different methods, and ii) identify and collect pure ColEVs devoid of other factors, such as ECs that often times co-purify with EVs^56^. Schematics for ColEVs isolation workflow are shown in **Figure 1**. The elution profiles of colonic digests from single bead without PVPP (-PVPP) is different from profile of gradient bead -PVPP (**Figure 2A**). The peaks labelled P1 for ColEVs and P2 for ColECs were collected and used for subsequent studies. Western blot assay for immunodetection of known markers of EVs showed that the isolated ColEVs were positive for the typical EV markers CD9 and CD63 while the P2 peak was negative (**Figure 2B**). The absence of the markers of EVs on the P2 (ECs) fraction is an indication that analytes in P2 may not be EVs. There were no differences in physical characteristics of ColEVs between single bead -PVPP and gradient bead -PVPP isolated ColEVs with regards to particle concentration (∼1.7e+10 particles/mL vs ∼2.7e+10 particles/mL), size (∼132.7 nm vs 136.6 nm), and ζ-potential (−22.4 mV vs −21.1 mV) respectively for single bead -PVPP and gradient bead -PVPP ColEVs (**Figure 2C**).

In the presence of PVPP (+PVPP), the elution profiles of colonic digests from are different (**Figure 2D**). The peaks labelled P1 for ColEVs and P2 for ColECs were collected and used for subsequent analyses. The use of PVPP did not affect the presence of CD9 and CD63 because both markers were present in P1 (ColEVs) while the P2 peak was negative (**Figure 2E**). The particle concentration (∼1.9e+10 particles/mL vs ∼2.4e+10 particles/mL) of single bead +PVPP and gradient bead +PVPP isolated ColEVs respectively were not different (**Figure 2F, left**). However, significant differences exist between single bead +PVPP and gradient bead +PVPP isolated ColEVs with regards to particle size (∼90.3 nm vs 133.5 nm, p=0.0106: Welch’s correction), and ζ-potential (−22.4 mV vs −21.1 mV, p=0.0363: Welch’s correction) respectively for single bead +PVPP and gradient bead +PVPP ColEVs (**Figure 2F**).

TEM analysis showed that EPs isolated with single and gradient bead columns without PVPP (-PVPP) contain ColEVs (**Figure 2G**) but not significant numbers of ColECs (**Figure 2H**). Similarly, single and gradient bead columns with PVPP (+PVPP) contain ColEVs (**Figure 2I**) but not significant numbers of ColECs (**Figure 2J**). The ColEVs are heterogenous in size, shape, and electron density irrespective of the use of PVPP or not. This said, ColEVs isolated with gradient bead +PVPP (**Figure 2I, bottom**) seem cleaner compared to other methods. This TEM analysis confirmed that colonic content contains EVs but not ECs and agrees with the western blot data presented in **Figures 2B, 2E**. While ECs are not enriched in colonic content, the data presented in **Figures 2C, 2F** show that the P2 analytes in **Figures 2A, 2D** have size, concentration, and ζ-potential data. This is not surprising because these may represent other particles in the extracellular space besides ColEVs and ColECs.

### PVPP facilitates isolation of pure ColEVs that are internalized by human T and epithelial cells

Internalization assays revealed that ColEVs labeled with lipophilic fluorescent membrane DiR deep red stain were taken up by TZM-GFP indicator cells irrespective of whether they were isolated without PVPP (-PVPP, **Figure 3A, 3B**) or with PVPP (+PVPP, **Figure 3C, 3D**). Visual assessment of the images showed reduced DAPI signal in cells treated with ColEVs isolated with single and gradient bead -PVPP (**Figure 3A**) compared to those treated with ColEVs isolated with single and gradient bead +PVPP (**Figure 3C**). This observation indicates that ColEVs isolated without PVPP treatment is toxic to cells. To confirm this observation, we counted the number of cells in wells treated with different concentrations of ColEVs isolated with single or gradient beads -PVPP (**Figure 3B**) compared to ColEVs isolated with single or gradient beads +PVPP (**Figure 3D**). We confirmed that in the absence of PVPP, ColEVs may be toxic to cells because TZM-GFP cells did not significantly increase from the starting seeding density of 10,000 cells (**Figure 3B**). In contrast, TZM-GFP cells treated with ColEVs isolated with single or gradient bead +PVPP significantly double from the starting seeding density of 10,000 cells to a mean of 20214 vs 26905 at 24 h; 50168 vs 70010 at 48 h; 60955 vs 68789 at 72 h; 61699 vs 66385 at 96 h respectively for single bead +PVPP vs gradient bead +PVPP (**Figure 3D**). These data reveal that the addition of PVPP treatment in ColEVs isolation protocol may be a necessary step in preserving target cell viability.

**Figure 3:**
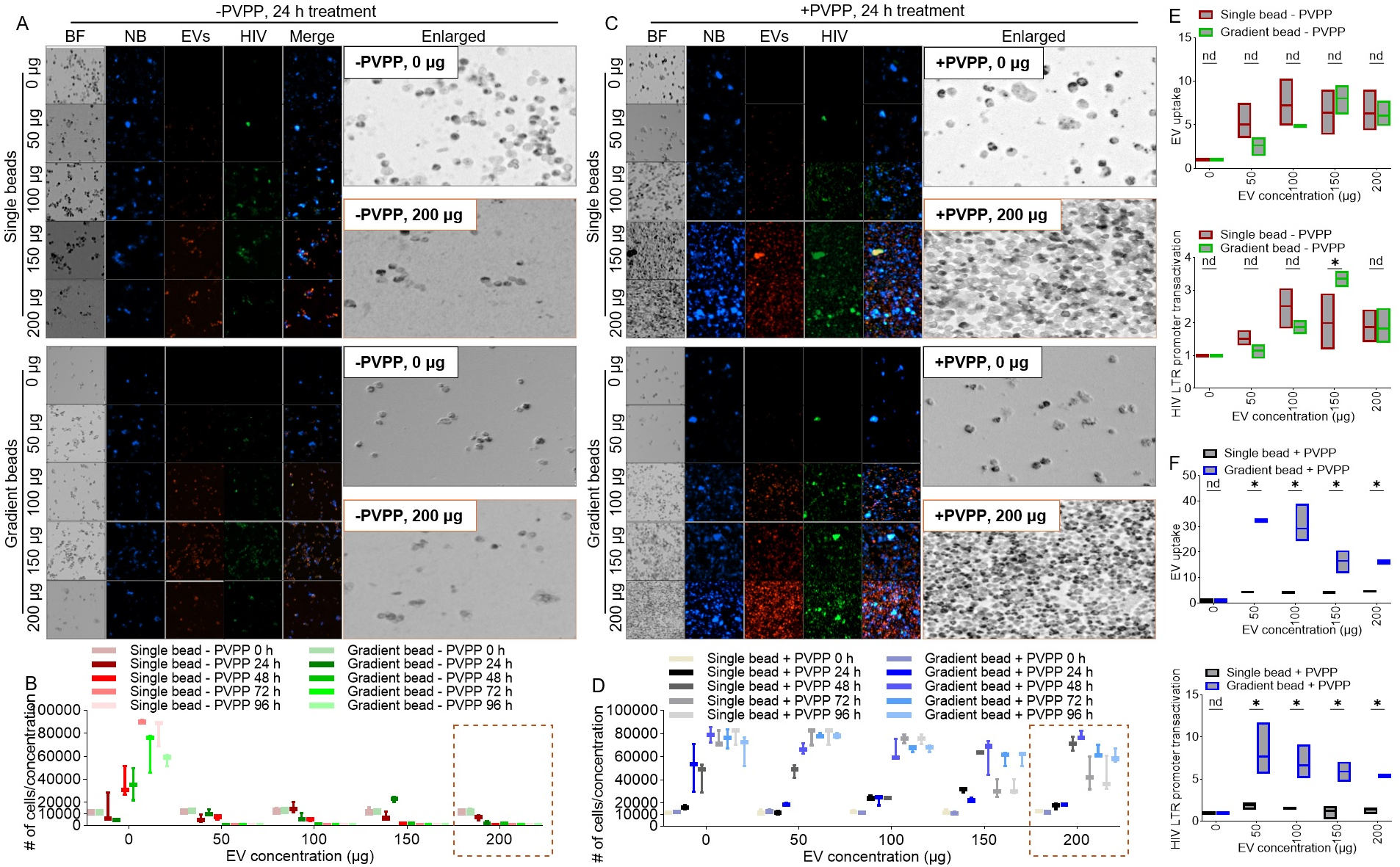
ColEVs are internalized by HIV latently infected JLAT-GFP T cells within 24 hours and the ColEVs transactivated HIV LTR promoter while the clarifying PVPP promotes survival of cells treated with ColEVs. A) Microscopic images of JLAT T cells treated with ColEVs isolated with single or gradient beads in the absence of PVPP. B) Quantification of JLAT T cells treated with ColEVs isolated with single or gradient beads in the absence of PVPP. C) Microscopic images of JLAT T cells treated with ColEVs isolated with single or gradient beads in the presence of PVPP. D) Quantification of JLAT T cells treated with ColEVs isolated with single or gradient beads in the presence of PVPP. E) Quantification of uptake (top) and level of HIV LTR promoter transactivation (bottom) of ColEVs isolated with single or gradient beads in the absence of PVPP and added to JLAT T cells. F) Quantification of uptake (top) and level of HIV LTR promoter transactivation (bottom) of ColEVs isolated with single or gradient beads in the presence of PVPP and added to JLAT T cells. Statistics were conducted with Prism Graph pad using Two-way ANOVA and two-stage linear step-up procedure of Benjamini, Krieger and Yekutieli. P*=<0.0001, ns=>0.9999. BF = bright field, NB = NucBlue.

Assessment of the uptake of ColEVs by cells revealed no difference in the uptake of single or gradient bead -PVPP isolated ColEVs (**Figure 3E, top**). However, in the presence of PVPP, internalization of ColEVs isolated with gradient beads was significantly higher compared to single bead isolated ColEVs (**Figure 3F, top**).

### ColEVs activate HIV LTR promoter

Internalization of ColEVs is suggestive of possible bioactivity. In this regard, single or gradient bead -PVPP isolated ColEVs had subtle effects on transactivating HIV LTR promoter (**Figure 3E, bottom**). In comparison, gradient bead +PVPP isolated ColEVs significantly increased HIV LTR promoter transactivation (**Figure 3F, bottom**). Both uptake of ColEVs and transactivation of HIV LTR promoter were assessed per cell. 50 µg of gradient bead +PVPP had the highest effect on transactivating HIV LTR promoter (**Figure 3F, bottom**).

### ColEVs isolated in the absence of PVPP interfere with MTT tetrazolium reduction

Visual assessment of cells treated with ColEVs, and enumeration of cell numbers (NucBlue staining and Brightfield) showed that ColEVs isolated in the absence of PVPP (-PVPP), irrespective of column type (single vs gradient bead) significantly decreased cell numbers (**Figure 3A**). In contrast, ColEVs isolated in the presence of PVPP (+PVPP) had no effect on cell numbers (**Figure 3B**), suggesting that in the absence of PVPP, ColEVs may have toxic effects on cells. Given the observations on cell numbers in **Figure 3**, we tested the effects of ColEVs on cell numbers (NucBlue staining) and cell viability using the MTT tetrazolium assay. While cell numbers decreased for wells seeded with cells in the presence of 100 µg ColEVs isolated without PVPP (**-PVPP, Figure 4A**, **Table 2**), MTT values increased in a concentration independent manner (**Figure 4B**, **Table 3**). In contrast, cells treated with 100 µg ColEVs isolated with PVPP increased in number (**Figure 4A**, **Table 2**) while MTT values did not change and was lower compared to the values from cells treated with ColEVs isolated without PVPP (**+PVPP, Figure 4B**, **Table 3**). These data suggest a direct interaction of ColEVs -PVPP with the MTT tetrazolium reduction.

**Figure 4:**
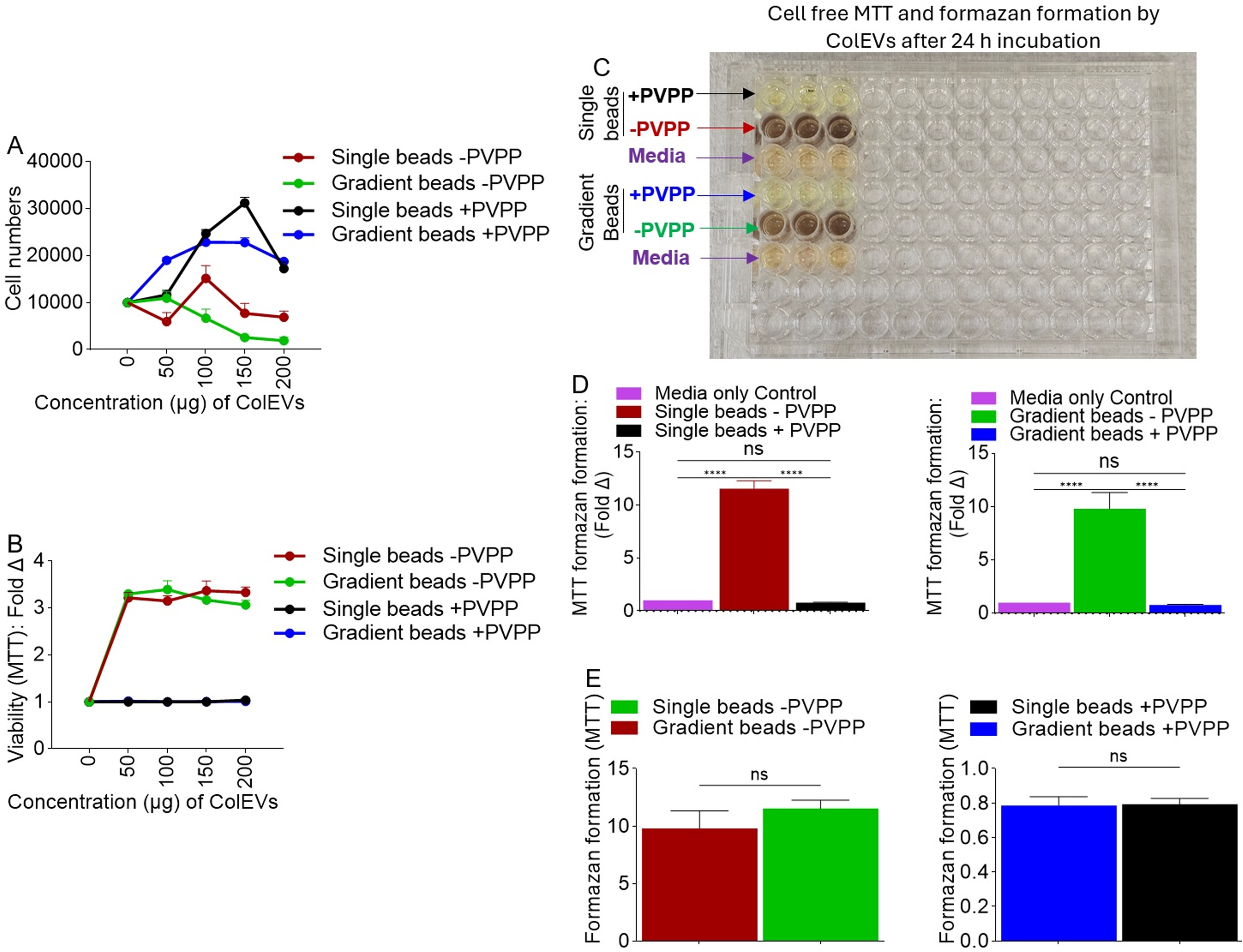
Effect of ColEVs on MTT reduction assay: A) The effect of ColEVs on cell number is dependent on PVPP and not on bead:column chemistry. B) The effect of ColEVs on cell viability is dependent on PVPP and not on bead:column chemistry. C) ColEVs isolated in the absence of PVPP irrespective of bead:column type caused very strong increase in formazan formation resulting in colorimetric changes in MTT reduction compared to ColEVs isolated in the presence of PVPP irrespective of bead:column type. Red and green arrows = wells with ColEVs –PVPP; black and blue arrows = wells with ColEVs +PVPP; black rectangles = wells with media alone. D, E) Quantification of MTT reduction by ColEVs isolated with single or gradient beads in the presence or absence of PVPP. Statistics were conducted with Prism GraphPad using unpaired t test with Welch’s correction. P****=<0.0001, P**=0.0026.

### Direct reductive potential of MTT absorbance by ColEVs -PVPP in a cell-free system

To confirm direct reductive potential of ColEVs, we used a cell-free system to assess the effects of ColEVs -PVPP and ColEVs +PVPP on tetrazolium reduction. Thus, ColEVs isolated with PVPP (+PVPP) or without PVPP (-PVPP) and with single or gradient columns were incubated with MTT and the absorbance was measured at 595 nm. Since color formation serves as a useful and convenient marker in the MTT assay, the results showed that wells containing ColEVs - PVPP had intense color change irrespective of whether the ColEVs were isolated with single vs gradient beads (**Figure 4C, red and green arrows**). In contrast, ColEVs +PVPP isolated with single and gradient did not change color irrespective of the of whether the ColEVs were isolated with single vs gradient beads (**Figure 4C, black and blue arrows**). The colors in wells containing ColEVs +PVPP (**Figure 4C, black and blue arrows**) were similar to the colors of the wells containing media alone (**Figure 4C, purple arrows**). The maximal absorbance values were significantly different, which were 0.79 (single bead, +PVPP) vs 11.5 (single bead, -PVPP) as shown in **Figure 4D**, **left**; and 0.79 (gradient bead, +PVPP) vs 9.79 (gradient bead, -PVPP) as shown in **Figure 4D**, **right.**

The reductive potential of MTT absorbance by ColEVs is dependent on the absence (**Figure 4E, left**) or presence (**Figure 4E, right**) of PVPP and not the PPLC column because MTT absorbance did not significantly change based on whether single or gradient bead column was used (**Figure 4E**). These results support interference in MTT data published by others^70,71^ and suggest that without the addition of PVPP, ColEVs reduce MTT in the absence of living cells.

Based on these data, the isolation method used for ColEVs can significantly influence the results of experiments using the MTT assay to measure the effects of EVs on cell growth. Subsequent studies will be conducted with ColEVs +PVPP isolated with single vs gradient beads to identify which isolation method is superior with regards to bioactivity of ColEVs.

### ColEVs isolated with single beads are more effective in transactivating HIV LTR promoter

JLAT-TAT-GFPcells were treated with DiR-labeled PBS (0 µg) or various concentrations (50, 100, 150, 200 µg) of ColEVs. Internalization of ColEVs and GFP expression as evidence of HIV LTR promoter reactivation were analyzed at 48 hours (**Figure 5A, top and bottom**), 72 hours (**Figure 5B, top and bottom**), and 96 hours (**Figure 5C, top and bottom**). Visual assessment of uptake of ColEVs (red florescence) and HIV LTR promoter activation (green fluorescence) are shown as representative microscopic images (**Figure 5D**). The uptake and bioactivity of ColEVs on HIV LTR promoter transactivation was confirmed in TZM-GFP cells at 96 hours by quantification of florescence intensities (**Figure 5E, top and bottom**) and the representative images are shown (**Figure 5F**). In both JLAT-TAT-GFP and TZM-GFP cells, ColEVs isolated with single beads significantly transactivated HIV LTR promoter at 72 and 96 hours post treatment (**Figures 5B, C, E, bottom panels**).

**Figure 5:**
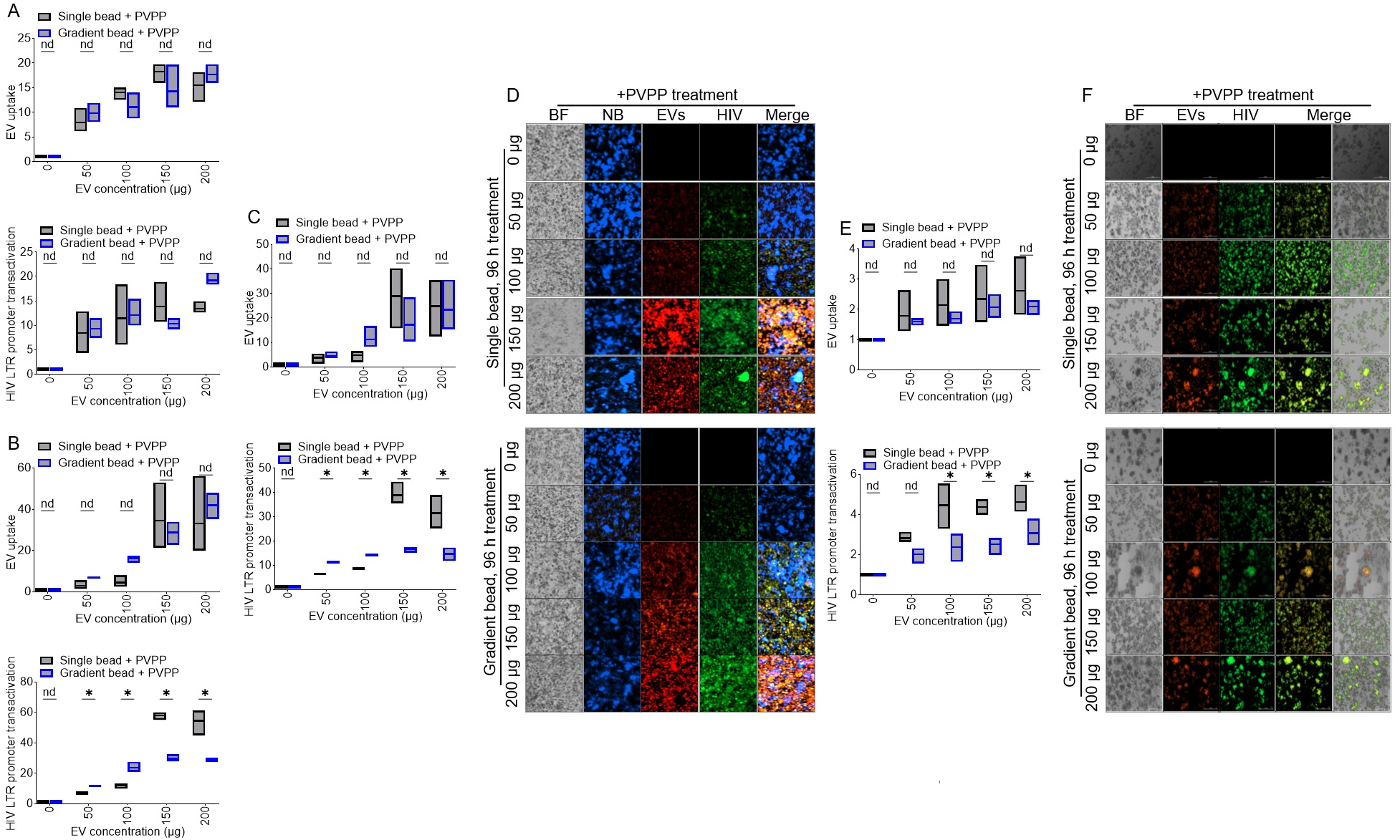
ColEVs isolated with single beads are more effective in transactivating HIV LTR promoter: **A**) Quantification of JLAT T cells response to ColEVs uptake (top) and level of HIV transactivation (bottom) following 48-hour treatment with ColEVs isolated with single or gradient beads in the presence of PVPP. **B**) Quantification of JLAT T cells response to ColEVs uptake (top) and level of HIV transactivation (bottom) following 72-hour treatment with ColEVs isolated with single or gradient beads in the presence of PVPP. **C**) Quantification of JLAT T cells response to ColEVs uptake (top) and level of HIV transactivation (bottom) following 96-hour treatment with ColEVs isolated with single or gradient beads in the presence of PVPP. **D**) Microscopic images of JLAT T cells response to ColEVs following 96-hour treatment with ColEVs isolated with single (top) or gradient (bottom) beads in the presence of PVPP. **E**) Quantification of TZM-GFP cells response to ColEVs uptake (top) and level of HIV transactivation (bottom) following 96-hour treatment with ColEVs isolated with single or gradient beads in the presence of PVPP. **F**) Microscopic images of TZM-GFP cells response to ColEVs following 96-hour treatment with ColEVs isolated with single (top) or gradient (bottom) beads in the presence of PVPP. Statistics were conducted with Prism GraphPad using the Two-stage linear step-up procedure of Benjamini, Krieger and Yekutieli. P*=<0.0001 to 0.0003, ns=>0.9999.

### ColEVs are originate from both bacteria and host particles

Here, we sought to identify the composition of ColEVs using flow cytometry (**Figure 6A**). To distinguish ColEVs of bacteria and host origins, we labelled ColEVs with antibodies against host CD9 and *Escherichia coli (E. coi)* AcrA protein (a component of the multi-drug efflux complex AcrAB-TolC that pumps out an extraordinarily wide variety of antibiotics, chemotherapeutic agents, detergents and dyes across two membranes). We found that both CD9+ and AcrA+ ColEVs are present in rhesus macaque GI. 46% of ColEVs are of host origin as they are CD9+, 52% are of bacteria origin as they are AcrA+, while 0.3% of ColEVs are hybrid particles because they are CD9+AcrA+ (**Figure 6B**).

**Figure 6:**
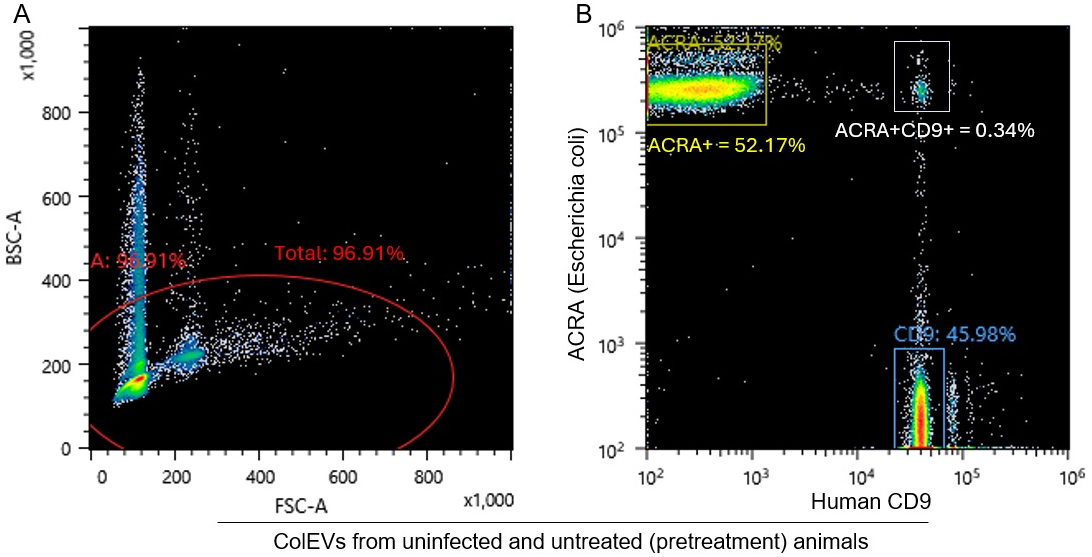
Flow cytometric analysis of the components of ColEVs: **A**) Gating strategy used to identify CD9+ and AcrA+ ColEVs. **B**) Plots of AcrA+, CD9+, and AcrA+CD9 ColEVs.

## Discussion

Before we finalized the efficient protocol described above, we tested different parameters – gradient bead column vs single bead column, isolation with PPVP vs isolation without PPVP. Thus, test samples were grouped into 4: single bead column with PPVP, Single bead column without PPVP, Gradient bead column with PPVP, and Gradient bead column without PPVP. We observed that the EV-rich peaks were more resolved among samples isolated with PPVP inclusive as compared to the samples that were isolated without the addition of PPVP (Figure 2). Additionally, the Transmission electron microscopy (TEM) images for samples that were isolated with PPVP were cleaner, with lesser non-EV fragments (**Figure 4C**), unlike the PPVP-devoid samples (**Figure 3C**). However, the concentration was higher in samples isolated with gradient beads + PPVP than in samples isolated with single beads + PPVP. EV-specific tetraspanin protein expression (**Figures 2B, 2E**) and TEM images (**Figure 2G to 2J**) confirmed the abundance of EVs in the isolates in peak 1 of the spectral profile and the absence of ColECs in the analytes. Overall, this study reveals that PPLC guarantees the isolation of a high yield purified EVs from colonic content, and that PVPP further improves the purity of the isolated ColEVs by binding to phenolic compounds and other contaminants that may interfere with the quality and quantity of the isolated products.

## Acknowledgments

This work was funded by NYMC/Lovelace Start-up funds, the National Institutes of Health (Grant # R01DA042348 (to CMO); Grant # R01DA042524, and R01DA052845 (to MM), P30AI161943, P51OD011104, P51OD111033. The work was supported in part by New York Medical College, Lovelace Biomedical Institute, and Texas Biomedical Research Institute core facilities.

## Author Contributions

MM-W oversaw the animal care and collected colon samples. NCJA wrote the protocol, isolated the EVs and validated the EV isolates. CMO prepared the figures that MM commented on. WN conducted data analysis. NCA, LSP, WN, MM, and CMO reviewed the draft for submission.

---

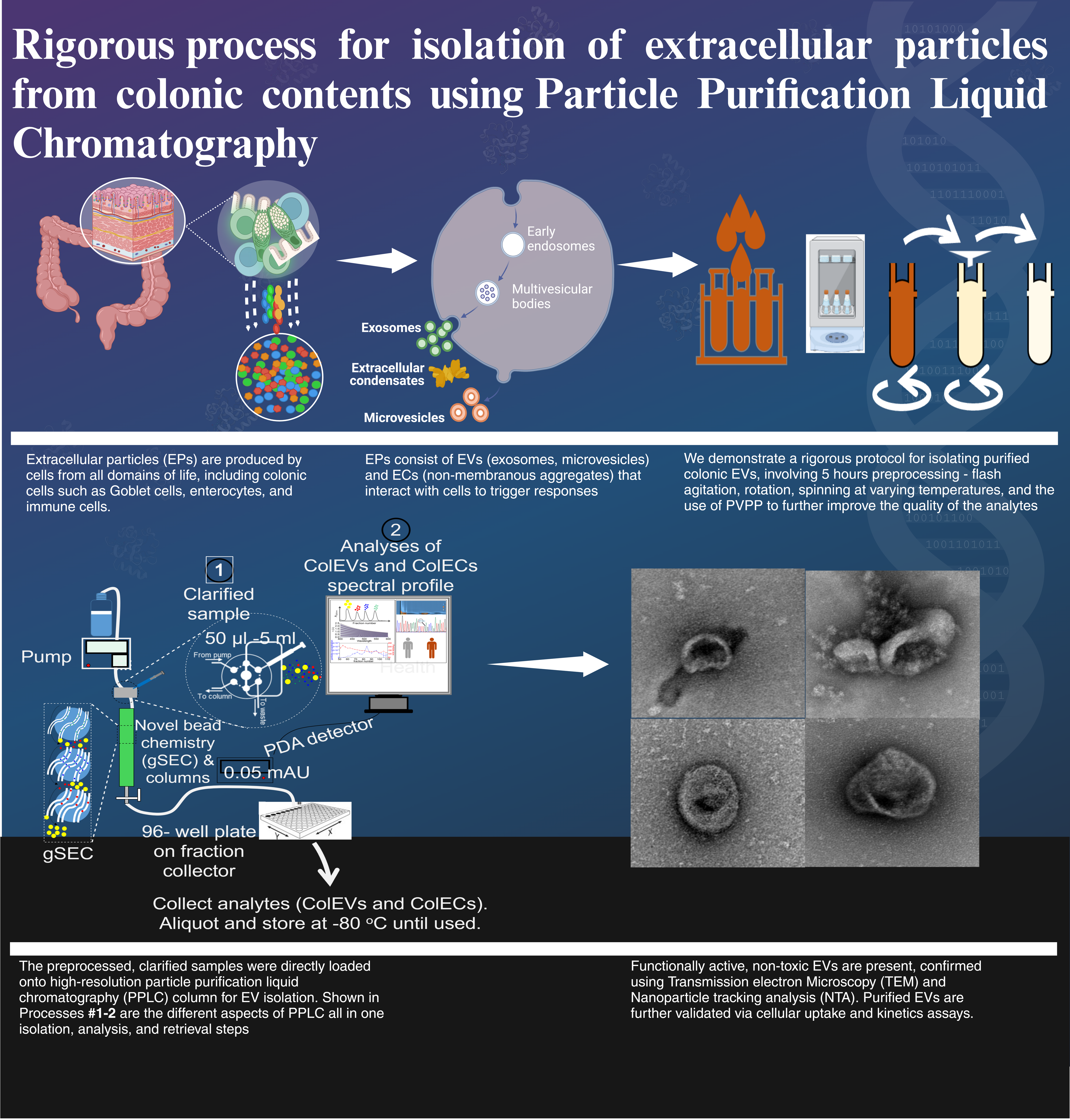

## References

1. Welch, J.L., Stapleton, J.T. & Okeoma, C.M. Vehicles of intercellular communication: exosomes and HIV-1. J Gen Virol (2019).

2. Admyre, C. et al. Exosomes with major histocompatibility complex class II and co-stimulatory molecules are present in human BAL fluid. Eur Respir J 22, 578–83 (2003).

3. Admyre, C. et al. Exosomes with immune modulatory features are present in human breast milk. J Immunol 179, 1969–78 (2007).

4. Baum, M.K. et al. Crack-cocaine use accelerates HIV disease progression in a cohort of HIV-positive drug users. J Acquir Immune Defic Syndr 50, 93–9 (2009).

5. Bobrie, A., Colombo, M., Raposo, G. & Thery, C. Exosome secretion: molecular mechanisms and roles in immune responses. Traffic 12, 1659–68 (2011).

6. Caby, M.P., Lankar, D., Vincendeau-Scherrer, C., Raposo, G. & Bonnerot, C. Exosomal-like vesicles are present in human blood plasma. Int Immunol 17, 879–87 (2005).

7. Lotvall, J. & Valadi, H. Cell to cell signalling via exosomes through esRNA. Cell Adh Migr 1, 156–8 (2007).

8. Madison, M.N., Roller, R.J. & Okeoma, C.M. Human semen contains exosomes with potent anti-HIV-1 activity. Retrovirology 11, 102 (2014).

9. Palanisamy, V. et al. Nanostructural and transcriptomic analyses of human saliva derived exosomes. PLoS One 5, e8577 (2010).

10. Pisitkun, T., Shen, R.F. & Knepper, M.A. Identification and proteomic profiling of exosomes in human urine. Proc Natl Acad Sci U S A 101, 13368–73 (2004).

11. Simons, M. & Raposo, G. Exosomes--vesicular carriers for intercellular communication. Curr Opin Cell Biol 21, 575–81 (2009).

12. Smith, J.A. & Daniel, R. Human vaginal fluid contains exosomes that have an inhibitory effect on an early step of the HIV-1 life cycle. Aids 30, 2611–2616 (2016).

13. Thery, C., Ostrowski, M. & Segura, E. Membrane vesicles as conveyors of immune responses. Nat Rev Immunol 9, 581–93 (2009).

14. Vojtech, L. et al. Exosomes in human semen carry a distinctive repertoire of small non-coding RNAs with potential regulatory functions. Nucleic Acids Res 42, 7290–304 (2014).

15. Kopcho, S., McDew-White, M., Naushad, W., Mohan, M. & Okeoma, C.M. SIV Infection Regulates Compartmentalization of Circulating Blood Plasma miRNAs within Extracellular Vesicles (EVs) and Extracellular Condensates (ECs) and Decreases EV-Associated miRNA-128. Viruses 15, 622 (2023).

16. Kopcho, S., McDew-White, M., Naushad, W., Mohan, M. & Okeoma, C.M. Alterations in Abundance and Compartmentalization of miRNAs in Blood Plasma Extracellular Vesicles and Extracellular Condensates during HIV/SIV Infection and Its Modulation by Antiretroviral Therapy (ART) and Delta-9-Tetrahydrocannabinol (Δ9-THC). Viruses 15, 623 (2023).

17. Kaddour, H. et al. Chronic delta-9-tetrahydrocannabinol (THC) treatment counteracts SIV-induced modulation of proinflammatory microRNA cargo in basal ganglia-derived extracellular vesicles. J Neuroinflammation 19, 225 (2022).

18. Chen, X.D. et al. Gut-Derived Exosomes Mediate Memory Impairment After Intestinal Ischemia/Reperfusion via Activating Microglia. Mol Neurobiol 58, 4828–4841 (2021).

19. Lyu, Y., Kopcho, S., Mohan, M. & Okeoma, C.M. Long-Term Low-Dose Delta-9-Tetrahydrocannbinol (THC) Administration to Simian Immunodeficiency Virus (SIV) Infected Rhesus Macaques Stimulates the Release of Bioactive Blood Extracellular Vesicles (EVs) that Induce Divergent Structural Adaptations and Signaling Cues. Cells 9(2020).

20. Ratajczak, J., Wysoczynski, M., Hayek, F., Janowska-Wieczorek, A. & Ratajczak, M.Z. Membrane-derived microvesicles: important and underappreciated mediators of cell-to-cell communication. Leukemia 20, 1487–95 (2006).

21. Valadi, H. et al. Exosome-mediated transfer of mRNAs and microRNAs is a novel mechanism of genetic exchange between cells. Nat Cell Biol 9, 654–9 (2007).

22. Schillaci, O. et al. Exosomes from metastatic cancer cells transfer amoeboid phenotype to non-metastatic cells and increase endothelial permeability: their emerging role in tumor heterogeneity. Sci Rep 7, 4711 (2017).

23. Madison, M.N., Jones, P.H. & Okeoma, C.M. Exosomes in human semen restrict HIV-1 transmission by vaginal cells and block intravaginal replication of LP-BM5 murine AIDS virus complex. Virology 482, 189–201 (2015).

24. Madison, M.N., Welch, J.L. & Okeoma, C.M. Isolation of Exosomes from Semen for in vitro Uptake and HIV-1 Infection Assays. Bio Protoc 7(2017).

25. Kopcho, S., McDew-White, M., Naushad, W., Mohan, M. & Okeoma, C.M. Alterations in Abundance and Compartmentalization of miRNAs in Blood Plasma Extracellular Vesicles and Extracellular Condensates during HIV/SIV Infection and Its Modulation by Antiretroviral Therapy (ART) and Delta-9-Tetrahydrocannabinol (Δ(9)-THC). Viruses 15(2023).

26. Kopcho, S., McDew-White, M., Naushad, W., Mohan, M. & Okeoma, C.M. SIV Infection Regulates Compartmentalization of Circulating Blood Plasma miRNAs within Extracellular Vesicles (EVs) and Extracellular Condensates (ECs) and Decreases EV-Associated miRNA-128. Viruses 15(2023).

27. Welch, J.L., Kaddour, H., Schlievert, P.M., Stapleton, J.T. & Okeoma, C.M. Semen exosomes promote transcriptional silencing of HIV-1 by disrupting NF-kB/Sp1/Tat circuitry. J Virol (2018).

28. Welch, J.L., Kaddour, H., Schlievert, P.M., Stapleton, J.T. & Okeoma, C.M. Semen Exosomes Promote Transcriptional Silencing of HIV-1 by Disrupting NF-κB/Sp1/Tat Circuitry. J Virol 92(2018).

29. Welch, J.L. et al. Semen Extracellular Vesicles From HIV-1-Infected Individuals Inhibit HIV-1 Replication In Vitro, and Extracellular Vesicles Carry Antiretroviral Drugs In Vivo. J Acquir Immune Defic Syndr 83, 90–98 (2020).

30. Dantas-Pereira, L., Menna-Barreto, R. & Lannes-Vieira, J. Extracellular Vesicles: Potential Role in Remote Signaling and Inflammation in Trypanosoma cruzi-Triggered Disease. Frontiers in Cell and Developmental Biology 9, 3574–3574 (2021).

31. Madeira, R.P. et al. New Biomarker in Chagas Disease: Extracellular Vesicles Isolated from Peripheral Blood in Chronic Chagas Disease Patients Modulate the Human Immune Response. Journal of Immunology Research 2021(2021).

32. Moreira, L.R., Serrano, F.R. & Osuna, A. Extracellular vesicles of Trypanosoma cruzi tissue-culture cell-derived trypomastigotes: Induction of physiological changes in non-parasitized culture cells. PLOS Neglected Tropical Diseases 13, e0007163–e0007163 (2019).

33. Torró, L.M.d.P., Moreira, L.R. & Osuna, A. Extracellular vesicles in chagas disease: A new passenger for an old disease. Frontiers in Microbiology 9(2018).

34. Fu, F., Jiang, W., Zhou, L. & Chen, Z. Circulating Exosomal miR-17-5p and miR-92a-3p Predict Pathologic Stage and Grade of Colorectal Cancer. Translational oncology 11, 221–232 (2018).

35. Goto, T. et al. An elevated expression of serum exosomal microRNA-191, -121, -451a of pancreatic neoplasm is considered to be efficient diagnostic marker. BMC cancer 18, 116–116 (2018).

36. Hurwitz, S.N. et al. Proteomic profiling of NCI-60 extracellular vesicles uncovers common protein cargo and cancer type-specific biomarkers. Oncotarget 7, 86999–87015 (2016).

37. Liu, Q. et al. Circulating exosomal microRNAs as prognostic biomarkers for non-small-cell lung cancer. Oncotarget 8, 13048–13058 (2017).

38. Wang, J. et al. Plasma exosomes as novel biomarker for the early diagnosis of gastric cancer. Cancer Biomarkers 21, 805–812 (2018).

39. Claridge, B., Lozano, J., Poh, Q.H. & Greening, D.W. Development of Extracellular Vesicle Therapeutics: Challenges, Considerations, and Opportunities. Frontiers in Cell and Developmental Biology 9(2021).

40. Oves, M. et al. Exosomes: A Paradigm in Drug Development against Cancer and Infectious Diseases. Journal of Nanomaterials 2018, 6895464–6895464 (2018).

41. Ratajczak, J. et al. Embryonic stem cell-derived microvesicles reprogram hematopoietic progenitors: evidence for horizontal transfer of mRNA and protein delivery. Leukemia 20, 847–856 (2006).

42. Tran, P.H.L. et al. Aptamer-guided extracellular vesicle theranostics in oncology. Theranostics 10, 3849–3866 (2020).

43. Xie, X. et al. Progress in the application of exosomes as therapeutic vectors in tumor-targeted therapy. Cytotherapy 21, 509–524 (2019).

44. Zhang, X. et al. Engineered Extracellular Vesicles for Cancer Therapy. Advanced Materials 33, 2005709–2005709 (2021).

45. Alhamwe, B.A. et al. Extracellular Vesicles and Asthma-More Than Just a Co-Existence. International journal of molecular sciences 22(2021).

46. Bartel, S. et al. Human airway epithelial extracellular vesicle miRNA signature is altered upon asthma development. Allergy: European Journal of Allergy and Clinical Immunology 75, 346–356 (2020).

47. Martin-Ventura, J.L., Roncal, C., Orbe, J. & Blanco-Colio, L.M. Role of Extracellular Vesicles as Potential Diagnostic and/or Therapeutic Biomarkers in Chronic Cardiovascular Diseases. Frontiers in Cell and Developmental Biology 10, 81–81 (2022).

48. Fu, H., Hu, D., Zhang, L. & Tang, P. Role of extracellular vesicles in rheumatoid arthritis. Molecular immunology 93, 125–132 (2018).

49. Mustonen, A.M. et al. Characterization of hyaluronan-coated extracellular vesicles in synovial fluid of patients with osteoarthritis and rheumatoid arthritis. BMC Musculoskeletal Disorders 22, 1–11 (2021).

50. Zhang, B., Zhao, M. & Lu, Q. Extracellular Vesicles in Rheumatoid Arthritis and Systemic Lupus Erythematosus: Functions and Applications. Frontiers in Immunology 11, 3451–3451 (2021).

51. Chun, H.J., Reis, R.L., Motta, A. & Khang, G. Biomimicked biomaterials : advances in tissue engineering and regenerative medicine. (2020).

52. Hao, Z.C. et al. Stem cell-derived exosomes: A promising strategy for fracture healing. Cell Proliferation 50(2017).

53. Zhou, Y. & Xiao, Y. The Development of Extracellular Vesicle-Integrated Biomaterials for Bone Regeneration. Advances in Experimental Medicine and Biology 1250, 97–108 (2020).

54. Han, N.D. et al. Microbial liberation of N-methylserotonin from orange fiber in gnotobiotic mice and humans. Cell 185, 2495–2509.e11 (2022).

55. Alvarez, F.A. et al. Blood plasma derived extracellular vesicles (BEVs): particle purification liquid chromatography (PPLC) and proteomic analysis reveals BEVs as a potential minimally invasive tool for predicting response to breast cancer treatment. Breast Cancer Res Treat 196, 423–437 (2022).

56. Kaddour, H., Lyu, Y., Shouman, N., Mohan, M. & Okeoma, C.M. Development of Novel High-Resolution Size-Guided Turbidimetry-Enabled Particle Purification Liquid Chromatography (PPLC): Extracellular Vesicles and Membraneless Condensates in Focus. Int J Mol Sci 21(2020).

57. Kaddour, H., Tranquille, M. & Okeoma, C.M. The Past, the Present, and the Future of the Size Exclusion Chromatography in Extracellular Vesicles Separation. Viruses 13(2021).

58. Gludish, D.W. et al. TZM-gfp cells: a tractable fluorescent tool for analysis of rare and early HIV-1 infection. Sci Rep 10, 19900 (2020).

59. Jordan, A., Bisgrove, D. & Verdin, E. HIV reproducibly establishes a latent infection after acute infection of T cells in vitro. Embo j 22, 1868–77 (2003).

60. Jordan, A., Defechereux, P. & Verdin, E. The site of HIV-1 integration in the human genome determines basal transcriptional activity and response to Tat transactivation. Embo j 20, 1726–38 (2001).

61. Willms, E., Cabañas, C., Mäger, I., Wood, M. & Vader, P. Extracellular vesicle heterogeneity: subpopulations, isolation techniques and diverse functions in cancer progression. Frontiers in immunology 9, 738 (2018).

62. Mathieu, M., Martin-Jaular, L., Lavieu, G. & Théry, C. Specificities of secretion and uptake of exosomes and other extracellular vesicles for cell-to-cell communication. Nature cell biology 21, 9 (2019).

63. Abdullahi, A.D. et al. Antibacterial activities of Miang extracts against selected pathogens and the potential of the tannin-free extracts in the growth inhibition of Streptococcus mutans. PLoS One 19, e0302717 (2024).

64. Ciferri, M.C., Quarto, R. & Tasso, R. Extracellular Vesicles as Biomarkers and Therapeutic Tools: From Pre-Clinical to Clinical Applications. Biology 10(2021).

65. Irmer, B., Chandrabalan, S., Maas, L., Bleckmann, A. & Menck, K. Extracellular Vesicles in Liquid Biopsies as Biomarkers for Solid Tumors. Cancers 15(2023).

66. Urabe, F. et al. Extracellular vesicles as biomarkers and therapeutic targets for cancer. American journal of physiology. Cell physiology 318, C29–C39 (2020).

67. Zhu, J., Wang, S., Yang, D., Xu, W. & Qian, H. Extracellular vesicles: emerging roles, biomarkers and therapeutic strategies in fibrotic diseases. Journal of Nanobiotechnology 2023 21:1 21, 1–19 (2023).

68. Kaddour, H. et al. Proteomics Profiling of Autologous Blood and Semen Exosomes from HIV-infected and Uninfected Individuals Reveals Compositional and Functional Variabilities. Mol Cell Proteomics 19, 78–100 (2020).

69. Chang, C.S. & Kao, C.Y. Current understanding of the gut microbiota shaping mechanisms. J Biomed Sci 26, 59 (2019).

70. Bruggisser, R., von Daeniken, K., Jundt, G., Schaffner, W. & Tullberg-Reinert, H. Interference of plant extracts, phytoestrogens and antioxidants with the MTT tetrazolium assay. Planta Med 68, 445–8 (2002).

71. Peng, L., Wang, B. & Ren, P. Reduction of MTT by flavonoids in the absence of cells. Colloids Surf B Biointerfaces 45, 108–11 (2005).

